# Immunological surveillance using gSG6-P1 biomarker reveals spatio-temporal dynamics of *Anopheles* exposure and gaps in malaria risk assessment

**DOI:** 10.1101/2025.02.15.638477

**Authors:** Manop Saeung, Natapong Jupatanakul, Niramon Jampeesri, Aneta Afelt, Theeraphap Chareonviriyaphap, Sylvie Manguin

## Abstract

Malaria risk assessment often relies on the entomological inoculation rate (EIR), which quantifies infectious bites per person over time. However, this approach does not account for human behavioral variability, limiting its effectiveness in accurately reflecting malaria risk at the population level. Recent studies have attempted to estimate mosquito-bite risk through the measurement of antibody responses to *Anopheles* salivary peptides. However, this approach has yet to be applied to evaluate malaria transmission risks along the Thailand-Cambodia border. This study aims to employ an immune biomarker for *Anopheles* salivary peptide (anti-gSG6-P1) to identify key risk factors associated with exposure to *Anopheles* mosquitoes in Sisaket Province, a region bordering Thailand and Cambodia. Blood samples from the same set of 184 participants were collected via finger prick during each season: rainy (August 2022), cool-dry (December 2022), and hot-dry (April 2023). Anti-gSG6-P1 antibody levels were quantified using enzyme-linked immunosorbent assay (ELISA). Factor Analysis of Mixed Data (FAMD) revealed that seasonality exerts the strongest influence on antibody levels. This pattern was likely driven by human activities, particularly the frequency of rubber tapping activity in the rubber plantation area where *Anopheles* species, especially *Anopheles dirus,* are present. While this study successfully identifies seasonality and human factors as critical influences on antibody responses, it also highlights gaps in understanding the kinetics of anti-salivary peptide responses to *Anopheles* bites, particularly in species with low salivary peptides sequence similarity to that of *Anopheles gambiae*. These findings emphasize the need for the development of new serological tools tailored to malaria vectors in the Greater Mekong Subregion to enhance malaria risk assessment and improve vector control strategies.

## 1. Introduction

Malaria, a disease transmitted by *Anopheles* mosquitoes, remains a significant public health concern, particularly in underprivileged regions as this disease is strongly linked to poverty [1]. The World Health Organization (WHO) estimates 226–263 million malaria cases and over 500,000 deaths annually between 2000 and 2023 [2]. To reduce the malaria burden, WHO has implemented malaria control programs targeting both vectors and parasites [3, 4], resulting in a notable decline in malaria incidence [5]. Following successful control efforts, low-transmission regions have transitioned to elimination programs as part of WHO’s malaria elimination framework, with vector control tailored to local environments and mosquito behavior [4].

The Greater Mekong Subregion (GMS), encompassing Cambodia, China (Yunnan Province), Laos, Myanmar, Thailand, and Vietnam, is one of the paradigms that enable the transition of malaria from the control phase to the elimination phase [6]. From 2000 to 2015, the region reported over 200,000 cases annually, peaking at more than 600,000 cases in 2012 [7–10]. Several efforts from malaria control programs contributed to a substantial reduction of cases in the region [9]. Then, in 2015, the control program in this region shifted to the malaria elimination phase, with a target to achieve the goal by 2030 [6]. The GMS has progressed toward malaria elimination with less than 100,000 cases reported in 2020 and 2021 [10]. However, in subsequent years, malaria cases rose to more than 150,000 in 2022 and 225,000 in 2023 [2]. This increase is mostly driven by Myanmar, which experienced a resurgence of indigenous malaria cases [2]. Malaria transmission persists even with widespread access to interventions such as insecticide-treated nets (ITN), and indoor residual spraying (IRS), often occurring in settings like forests where these existing measures are less practical [11, 12]. Outdoor transmission, driven by the behaviors of forest workers and rubber tappers, highlights the complexity of residual transmission dynamics [11, 13].

Measuring malaria transmission risk is challenging, with variations in individual exposure within the same community. The entomological inoculation rate (EIR), which quantifies the number of infected mosquito bites per person over a specified period of time, is a standard measure but is subject to limitations, including site representativeness and collection biases. In addition, malaria vectors in this region predominantly inhabit forested areas [14], further complicating entomological surveys. Moreover, human vulnerability varies based on occupation, locality, and socioeconomic status [15, 16], highlighting the need for alternative tools to more accurately estimate malaria transmission risk.

One promising approach is using human immune responses as biomarkers for *Anopheles* bites. The gSG6-P1 peptide, derived from a conserved salivary protein in *Anopheles gambiae,* has been validated in malaria-endemic areas like Senegal, demonstrating a positive correlation between IgG antibody levels and mosquito exposure [17]. Previous studies reported a positive association between the gSG6-P1 peptide and exposure to bites of *An. gambiae*, suggesting its utility as a proxy for mosquito bites [18–22]. In Thailand, this peptide technique was validated in Tak Province, an area with highest recorded malaria incident where *Anopheles minimus* was the dominant species and found the potential use of this peptide as a biomarker in the GMS [23]. However, further validation is required in other areas with different dominant *Anopheles* species.

Sisaket Province was identified as one of the most significant malaria-endemic provinces along the Thai-Cambodia border, with *Anopheles dirus*, an exophagic and exophilic mosquito, as the predominant vector [14]. Although malaria control efforts have significantly reduced case numbers and advanced progress toward elimination, the region remains at risk of re-establishment due to the continued presence of vectors and vulnerability of people [14, 24].

Malaria transmission in Sisaket is primarily concentrated in rubber-forest areas―where rubber plantations border forests ―along the Thai-Cambodian border [24]. Our previous entomological study demonstrated seasonal variations of mosquito biting rates which peak during the rainy and cool-dry seasons [14]. These variations in mosquito prevalence and biting behaviors emphasize the necessity for tools to accurately assess malaria transmission risk at the population levels.

This study aims to validate the gSG6-P1 peptide as a biomarker for *Anopheles* bites among at-risk populations (rubber tappers) in Sisaket Province, focusing on its effectiveness in estimating mosquito exposure. By identifying key environmental and behavioral factors influencing biting rates, the study seeks to improve our understanding of malaria transmission dynamics and establish a reliable serological tool to monitor transmission risks in endemic regions.

## 2. Materials and Methods

### 2.1 Study locations

Three villages, Huai Chan, Kan Throm Tai, and Non Thong Lang, located in Khun Han District, Sisaket Province were chosen due to the histologically high malaria burden (***Figure 1***). These villages are gradually slope up toward the southern part, where large rubber plantations and primary forests are located. Kan Throm Tai and Non Thong Lang are situated in lower areas (∼200 m), surrounded by paddy fields and scattered rubber plantations. In contrast, Huai Chan Village is located at a higher altitude (∼240 m) and is mostly surrounded by rubber plantations, croplands, and deciduous forests. The southernmost area of Sisaket Province shares a border with Cambodia (***Figure 1***). Agriculture is the main occupation of villagers in these areas. Weather conditions are categorized into three seasons, which are rainy (May– October), cool-dry (October–February), and hot-dry (February–May) [25]. Spatial data was obtained from USAID website (https://landcovermapping.org/en/landcover/), processed using QGIS software (version 3.22.7).

**Figure 1.**
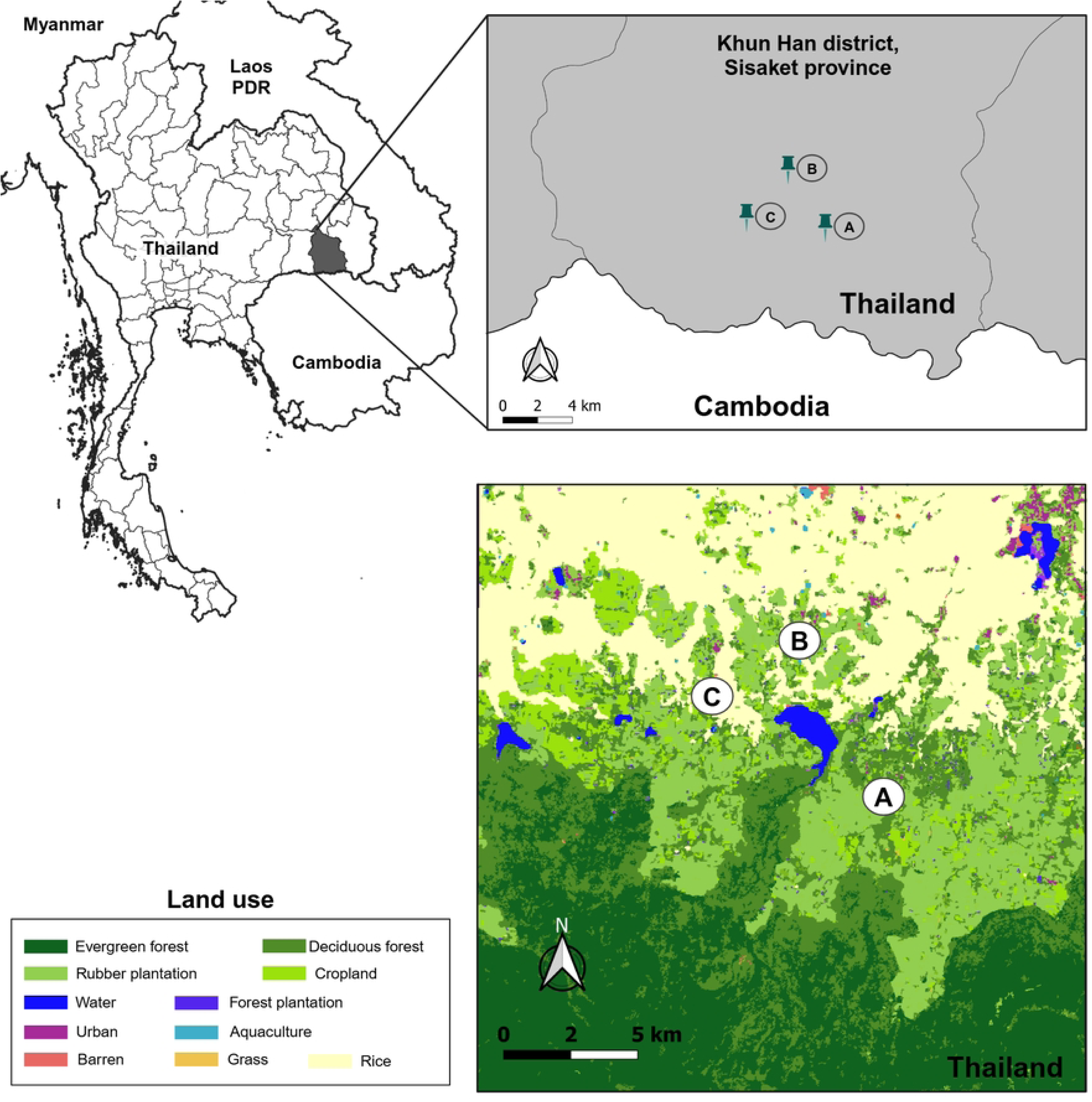
The study locations in Khun Han District, Sisaket Province, Northeastern Thailand. Three villages were selected, including Huai Chan (A), Kan Throm Tai (B), and Non Thong Lang (**C**).

### 2.2 Study design and inclusion criteria

A one-year cross-sectional survey was designed to capture the temporal dynamics of the anti-gSG6 P-1 peptide from local people in at-risk groups. Finger prick blood samples were collected one time/season, which started in rainy (August 2022), cool-dry (December 2022), and hot-dry seasons (April 2023). Participants were selected based on the following criteria: 1) age between 18–60 years old, 2) residing in these villages, 3) Thai citizenship, and 4) work in rubber plot. Each participant was followed up throughout the three seasons. A participant was considered a dropout if they were unable to provide a blood sample during either season.

### 2.3 Dry blood spot collection and interview

Finger prick blood was collected by local health authorities using Whatman No. 1 filter papers (Whatman International Ltd., Banbury, UK), size 3 × 12 cm following an approved protocol. Two blood spots were collected from each participant and labeled with a unique identification code. Each blood sample was dried before being kept in a plastic zip-lock bag to prevent contamination. All blood samples were then shipped to the National Center for Genetic Engineering and Biotechnology (BIOTEC) and stored at −20°C until further analysis.

To gather population characteristic data for analyzing factors influencing anti-salivary responses, information on participants’ engagement in rubber plantations was collected. Since seasonal variations influenced participation behavioral patterns, during each blood collection, participants were asked to report their average frequency of rubber plantation visits per week. Additionally, as the distance between participants’ homes and rubber plantations varied, they were asked to estimate this distance to assess potential spatial influences on exposure risk.

### 2.4 Antibody detection by enzyme-linked immunosorbent assay (ELISA)

The immunological assay to detect human antibody against *Anopheles* salivary peptide used in this study was adapted from a previous protocol [23]. A dry blood spot sample was cut into a circle shape (diameter: 6 mm) and eluted in 250 µL of 1X phosphate-buffered saline (1X PBS) with 0.1% Tween (Thermo Fisher Scientific, Cat# BP337-100) at 4°C for 48 hrs. Then, the filter paper was removed and eluted blood was divided into three tubes (∼80 µL/tube) before use to prevent degradation from the freeze-thaw process. All samples were kept at −20°C for further analysis with ELISA.

The assay was conducted in a 96-well ELISA plate (Life Sciences, Cat#3590) and began with the coating step. In this step, 100 µL of gSG6-P1 peptide (sequence: H-EKVWVDRDNVYCGHLDCTRVATF-OH) (0.2 µg/well diluted in 1X PBS) (GENEPEP) was added to two of 96-well per sample, while 100 µL of 0.1X blocking buffer in 1X PBS (Thermo Fisher Scientific, Cat# 37570) was added to another well as a negative control. The plate was incubated at 37°C for 3 hrs in an incubator (Vision Scientific, VS-1203P3L) and washed three times with 200 µL/well of washing solution (0.1% Tween in distilled water). To prevent nonspecific antibody binding to the remaining surface, 200 µL of 0.1X blocking buffer (diluted in 1X PBS) was added to each well. The plate was incubated at 37°C for 3 hrs and washed three times with 200 µL/well of washing solution. Next,100 µL of eluted blood sample (diluted 1:10 in 1X PBS & 1% Tween) was added to each well, allowing the antibodies to bind to the gSG6-P1 peptide. The plate was incubated overnight at 4°C followed by three washes with 200 µL washing solution. After washing, 100 µL of secondary antibody solution (0.05 µg/well diluted in 1X PBS & 1% Tween) (Biotin-conjugated Mouse Anti-human IgG, BD Biosciences, Cat# 555785) was added to each well and incubated at 37°C for 1 hr and 30 mins followed by three washes with 200 µL/well of washing solution. Following the washing steps, 100 µL of horse radish peroxidase-conjugated streptavidin solution (0.002 µg/well diluted in 1X PBS & 1% Tween) (Jackson ImmunoResearch Laboratories, Cat# 016-303-084) was added to each well and incubated at 37°C for 1 hr. After a final washing step, 100 µL of substrate solution (0.1 mg/well) containing 2,2’-azino-bis (3-ethylbenzothiazoline-6-sulfonic acid) (ABTS) (Sigma-Aldrich, Cat# A9941-100TAB) (diluted in 0.05M citrate buffer (pH4) & hydrogen peroxide (H_2_O_2_)) was added to each well. The plate was wrapped in aluminum foil and incubated at room temperature for 1 hr. The optical density (OD) was measured at 410 nm using an ELISA plate reader (BioTek, Synergy H1).

The absolute antibody levels obtained from each well were corrected by subtracting the noise value (negative control). The corrected values were expressed as delta optical density (ΔOD) and used to calculate the average antibody level for each sample, which was performed in duplicate. To account for plate-to-plate variations, normalization of ΔOD values was performed using standard curves generated from five blood samples representing different levels of antibody responses. This approach ensured consistency and reliability across all assay plates.

### 2.5 Statistical analysis and visualizations

The adjusted OD values from all ELISA samples were analyzed to identify quantitative and qualitative factors influencing variability of anti-salivary peptide responses using Factor Analysis for Mixed Data (FAMD) using FactoMineR package [26] in RStudio (Version 4.2.1). FAMD is a variant of Multiple Factor Analysis that integrates principal component analysis for continuous variables and multiple correspondence analysis for categorical variables, allowing for the simultaneous analysis of both data types [27].

For this analysis, three categorical variables (season, village, and distance from house to rubber plot) and two continuous variables (frequency of entering rubber plot per week (RubberPerWeek), and age of volunteers) were included. Prior to analysis, antibody levels were ranked in ascending order and classified into five groups based on the number of data points, with the average antibody level calculated for each group. FAMD was used to visualize variables in a multi-dimensional space, capturing their interactions. The first two dimensions, which explained the majority of data variance, were selected for further analysis. Variables within each dimension were assessed based on their contribution values, with those exceeding the expected average considered significant contributors to the principal dimensions. A FAMD factor map was then plotted to observe patterns in antibody levels and identify key influencing variables.

Statistical comparisons of antibody levels between two independent groups was performed using the Wilcoxon Rank-Sum Test, while comparisons between two dependent groups were conducted using the Wilcoxon Signed-Rank Test. Statistical significance was defined as a p-value <0.05. The data were analyzed using jamovi 2.3.6 (jamovi.org) and data visualization was performed in RStudio program (Version 4.2.1).

## 3. Results

### 3.1 Population characteristics

A total of 184 participants who reside in one of three villages, aged between 18–60 years old, and provided blood in every season were included in data analyses. Among the three villages, a majority of participants were from Kan Throm Tai (52.72%), followed by Huai Chan (32.07%), and Non Thong Lang (15.22%) villages. Females constituted a larger proportion (63.05%) compared to males (36.95%). Participants were categorized into ten-year age intervals within the population aged 18–60 years, with the majority of participants (67%) falling into two working age groups: 41–50 years (34%) and 51–60 years (33%). The distance between rubber plot and participants’ homes was classified into three groups: <1 km, 1–5 km, and >5 km. Most participants (55.43%) live > lived more than 5 km away from the rubber plot 41.30% residing within 1–5 km, and only 3.26% living less than 1 km away from the rubber plot. Rubber plot entry activity varied by season, with a majority reporting a higher frequency (5–7 day/week) during the cool-dry (104, 56.5%) and hot-dry (99, 53.8%) seasons compared to the rainy season (55, 29.9%) (***Table 1***).

**Table 1.**
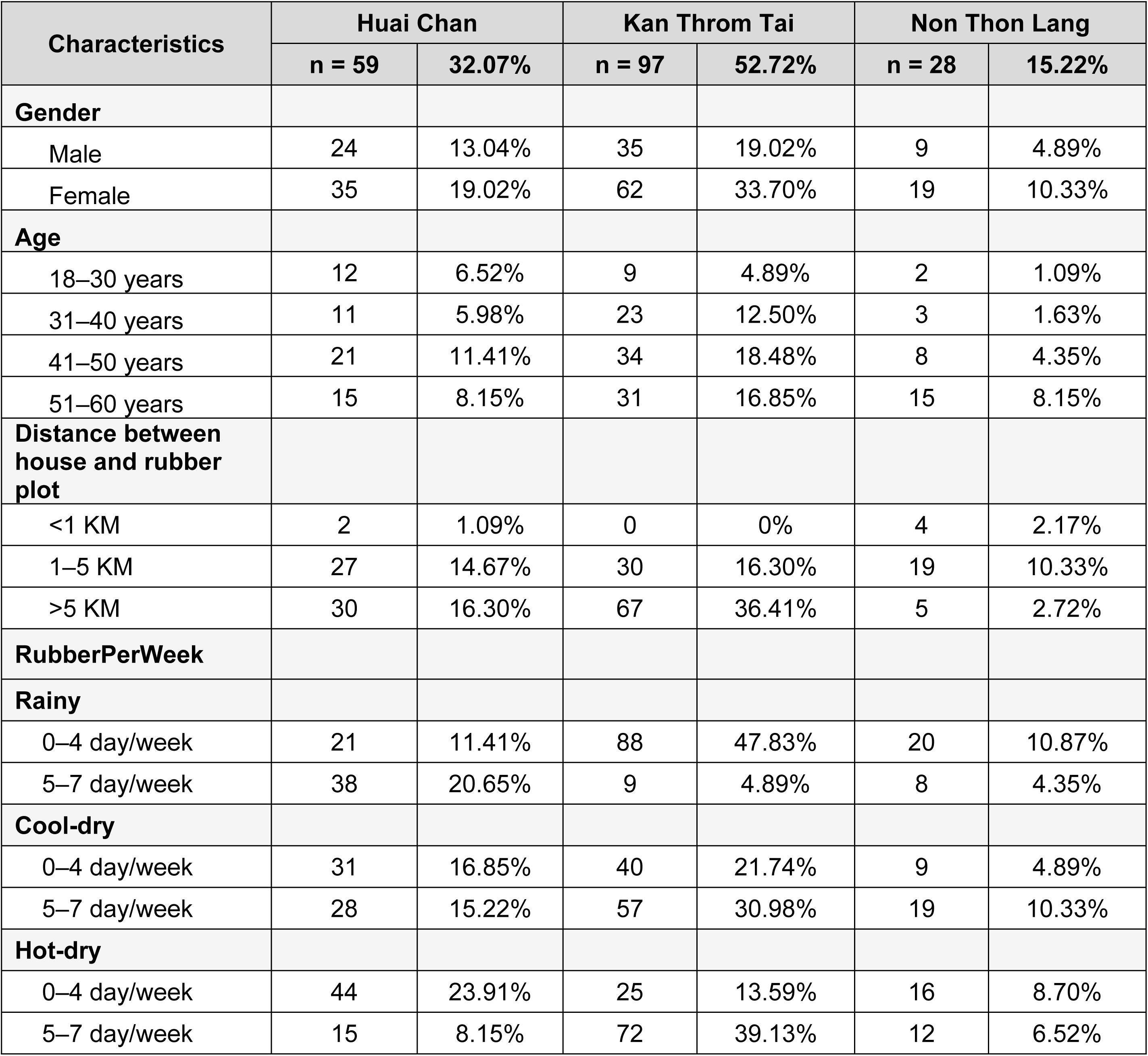
Characteristics of the 184 participants from Sisaket Province (age 18*–*60 years old), from August 2022 to April 2023.

### 3.2 Exploring key variables influencing antibody responses using FAMD

To identify the key factors influencing antibody levels, we conducted FAMD to explore relationships between population characteristics and antibody levels. The first step of FAMD was to calculate the principal dimensions (Dims), which combine variances of the original factors including: season, antibody level, frequency entering rubber plot/week, age, village, and distance from house to rubber plot, helping to simplify the relationship between factors in the dataset. Scree plot (***Figure 2A***) were used to illustrate the proportion of total variance in the dataset explained by each Dim. Dims 1 and 2 accounted for the largest share, together explaining 30.76% of the total variance. This indicates a high level of complexity in the dataset and suggests that multiple factors contribute to the observed variations in antibody levels.

**Figure 2.**
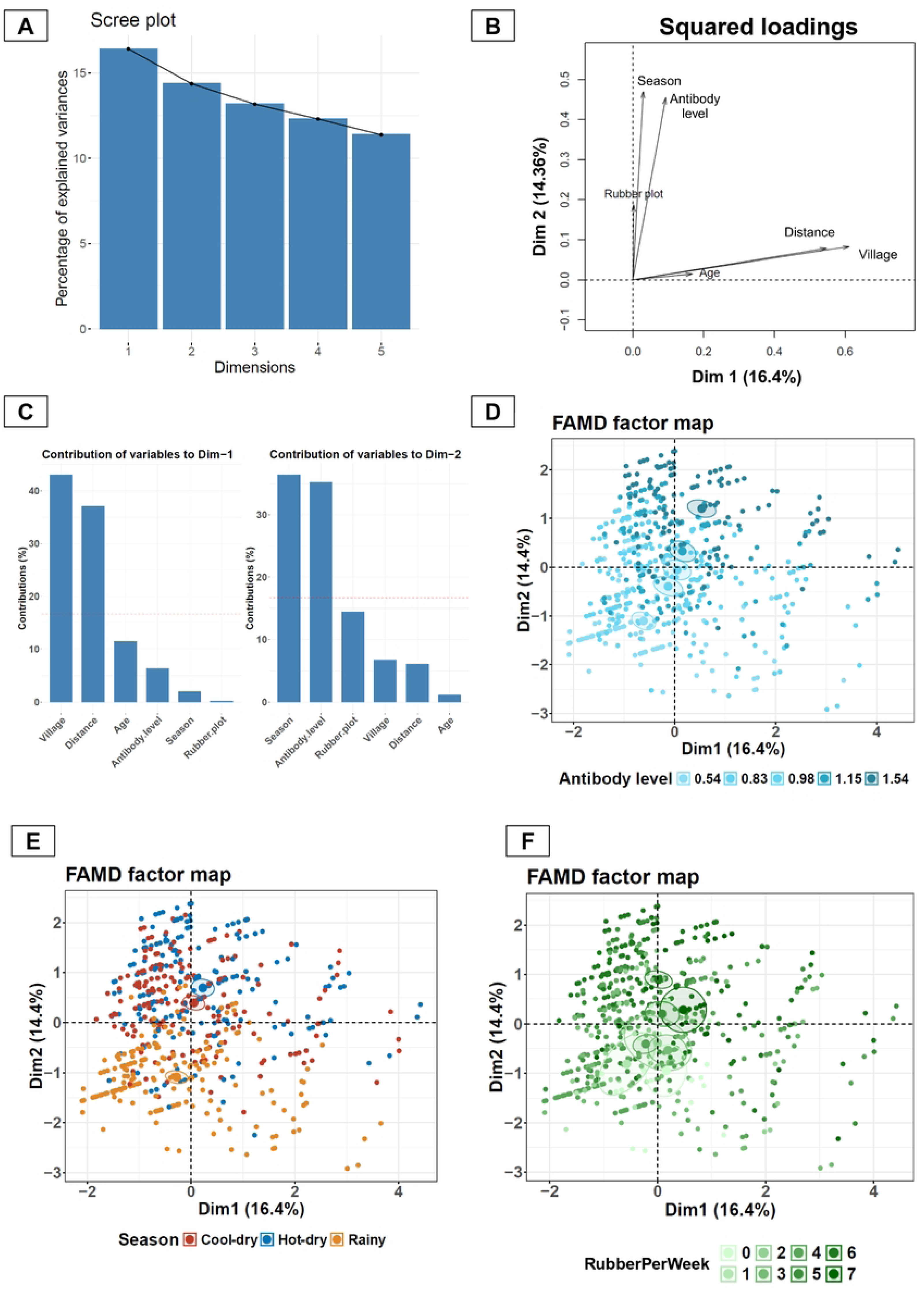
FAMD analysis reveals underlying variables influencing the level of anti-gSP6 P1 responses. (**A**) Scree plot demonstrating the contribution of each dimension to the total variance of the dataset. (**B**) Squared loadings plot demonstrating the correlations between categorical and continuous variables with Dims 1 and 2. (**C**) Contribution plot demonstrating contributions of each variable to Dims 1 and 2. (**D-F**) FAMD factor maps represents coordinate of each sample on the Dims 1 and 2, with color overlays indicating **(D)** antibody response levels, (**E**) season, and (**F**) frequency of rubber plot entry per week (RubberPerWeek).

Key determinants influencing antibody levels were identified using the squared loading plot and the percentage contribution of variables to Dim-1 and Dim-2 (***Figure 2B***). The results from these analyses revealed a correlation between anti-gSG6-P1 levels and both Dim-1 and Dim-2, with a stronger association observed with Dim-2. For Dim-2, the primary contributors were season (36%) and the frequency of entering rubber plot per week (RubberPerWeek; 35%), indicating that these factors are critical determinants of anti-gSG6-P1 levels (***Figure 2C***). Dim-1 had a lesser influence on anti-gSG6-P1 levels and the major contributors to this dimension were *village* (43%) and *distance from house to rubber plot* (37%) (***Figure 2C***).

Based on these findings, three key variables—antibody levels, season, and RubberPerWeek— were selected for further analysis to assess their directional relationships. The patterns of these variables were visualized using an FAMD factor map, illustrating their interactions within individual data points. The factor map of antibody levels reveals a gradient, with low-antibody level clusters located in the (-, -) quadrant (lower left) and high-level clusters in the (+, +) quadrant (upper right) (***Figure 2D***). The factor map for season showed that the rainy season corresponded to lower antibody levels, while the cool-dry and hot-dry seasons were associated with higher antibody levels (***Figure 2E*)**. Furthermore, the higher antibody levels observed during the cool-dry and hot-dry seasons were correlated with increased entries into rubber plots, as depicted in the factor map for RubberPerWeek (***Figure 2F***). Conversely, lower antibody levels during the rainy season were associated with reduced RubberPerWeek frequencies. These findings underline a strong relationship between seasonal rubber plot activity and antibody responses (***Figure 2F***).

### 3.3 Effect of rubber plot entry frequency on anti-gSP6 P1 levels

While FAMD identified rubber plot entry frequency as a factor correlated with anti-gSG6 P1 levels, it did not provide a direct statistical comparison between these variables. Based on the FAMD map, which shows clustering of participants with 0–4 days/week and 5–7 days/week of entry frequencies, we categorized participants into two groups: low frequency (0–4 days/week) and high frequency (5–7 days/week) then compared antibody responses between these two groups. The results indicated that individuals with a high frequency of rubber plot entry had significantly higher antibody responses (1.08 ± 0.36) compared to those with a low frequency (0.96 ± 0.35) (***Figure 3A***).

**Figure 3.**
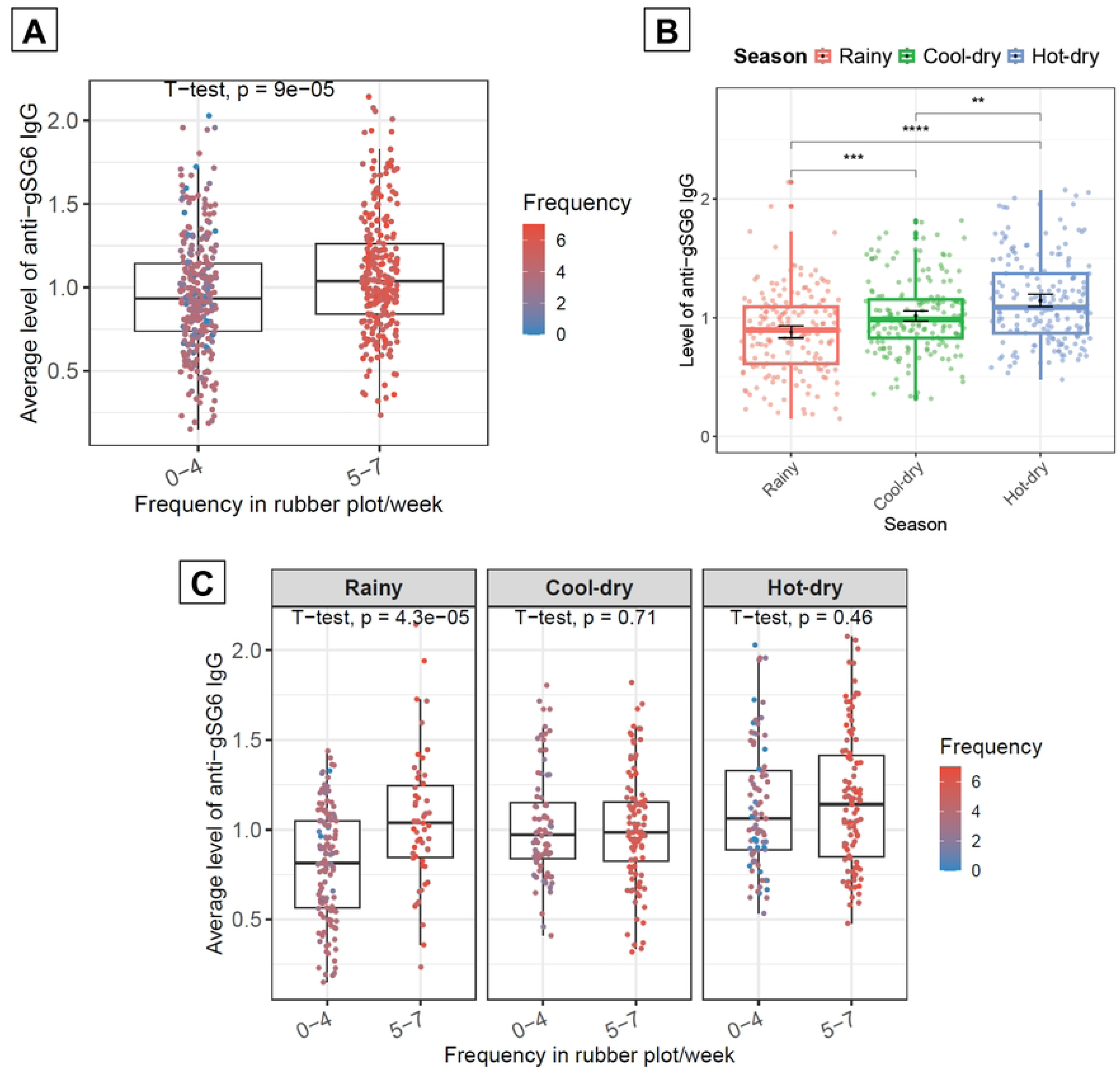
Frequency of entering the rubber plot/week and seasonal variables affect anti-gSG6 P1 response. (**A**) Box plot comparing levels of anti-gSG6 P1 between participants with low- and high-rubber plot visit frequency. Higher antibody responses were associated with more frequent visits to the rubber plot, regardless of seasonal variation. (**B**) Box plot demonstrating temporal dynamics of antibody responses across three seasons. (**C**) Box plot demonstrating effects the combined effect of season and rubber plot entry frequency on antibody levels. The box plot displays the interquartile range, and the whiskers indicate the range of maximum and minimum values. The dot plots display the raw data from each individual. Statistical analyses were performed using the Wilcoxon Rank-Sum Test, with significance levels indicated as *p*<0.01 (**) and *p*<0.001 (***), *p*<0.0001 (****), while NS indicates no statistically significant *p*>0.05.

To further explore the temporal dynamics of antibody levels, seasonal variations in anti-gSG6 P1 levels were analyzed. Antibody levels were lowest during the rainy season (0.88 ± 0.34), increased during the cool-dry season (1.02 ± 0.31), and peaked during the hot-dry season (1.15 ± 0.36) (***Figure 3B***). When considering both entry frequency and seasonal variation together, we observed distinct antibody level patterns between the high- and low-frequency groups across different seasons. During the rainy season, individuals with a high frequency of rubber plot entry had significantly elevated antibody levels (1.05 ± 0.37) compared to those with low entry frequency (0.81 ± 0.31). In contrast, during the cool-dry season, antibody levels in the low-frequency group increased to 1.03 ± 0.30, reaching a level comparable to the high-frequency group, which remained unchanged at 1.01 ± 0.31. During the hot-dry season, antibody levels were elevated across both groups, with high-frequency individuals reaching 1.16 ± 0.38 and low-frequency individuals at 1.12 ± 0.35 (***Figure 3C***).

### 3.4 Spatio-temporal dynamics of anti-gSG6 P1 antibody responses in the study population

Next, we investigated the spatio-temporal dynamics of antibody responses among residents of three villages over one year to assess potential variations in antibody levels across different locations. Our analysis revealed similar patterns in two villages, Kan Throm Tai and Non Thong Lang. In both villages, antibody levels were lowest during the rainy season (0.73 ± 0.31 and 0.83 ± 0.24, respectively) and increased during the cool-dry season (1.08 ± 0.29 and 1.02 ± 0.28) and hot-dry season (1.15 ± 0.38 and 1.21 ± 0.25, respectively) (***Figure 4***). In contrast, Huai Chan village had a distinct population antibody response pattern. Antibody levels were highest during the rainy season (1.15 ± 0.27), decreased to their lowest during the cool-dry season (0.91 ± 0.32), and then increased again during the hot-dry season (1.10 ± 0.38) (***Figure 4***). This unique pattern of elevated antibody levels observed in Huai Chan during the rainy season may be due to a higher proportion of residents in Huai Chan frequently entering rubber plots during rainy season (5–7 times per week) compared to the other two villages (***Table 1***).

**Figure 4.**
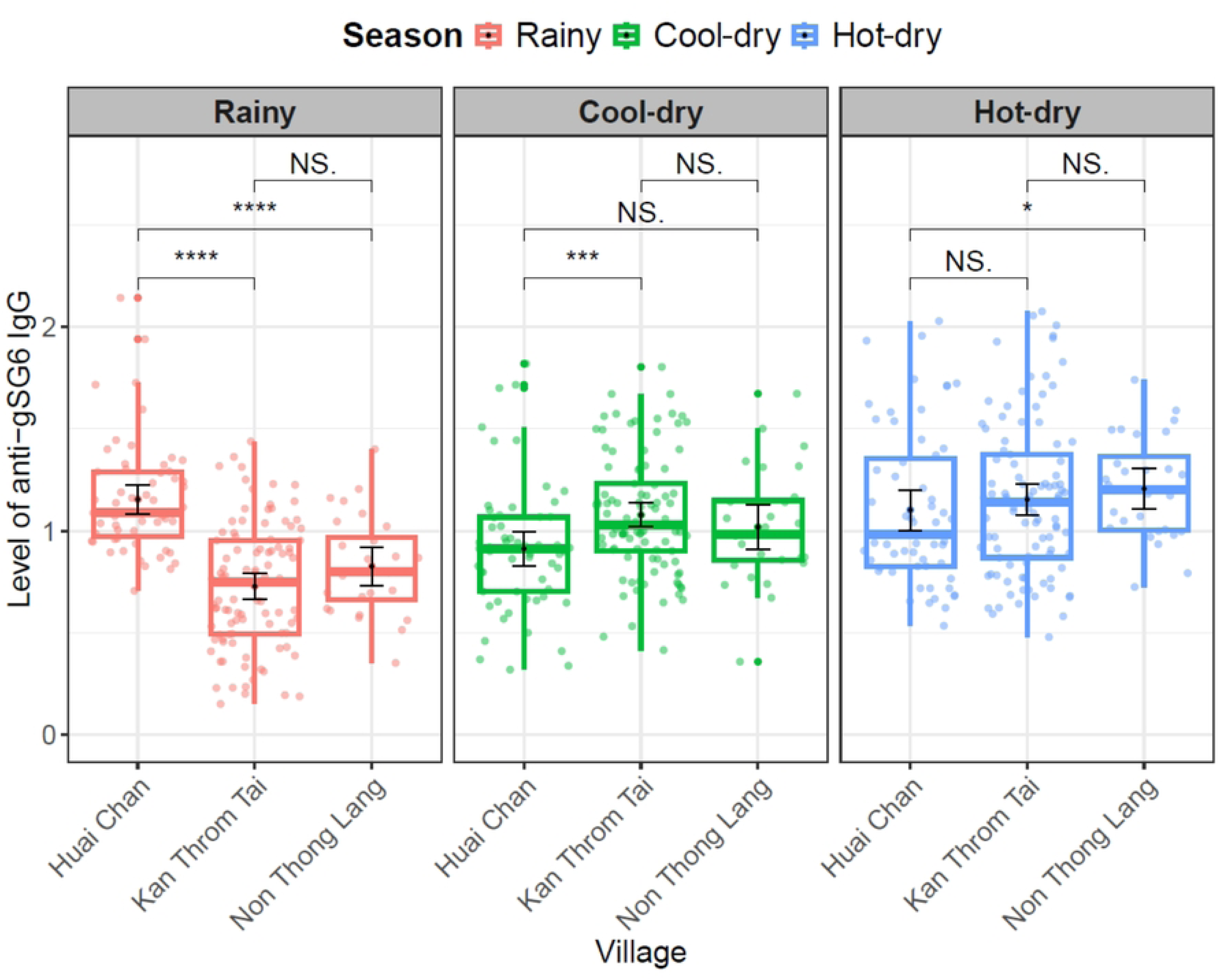
Spatial-temporal dynamics of antibody responses among all participants across three seasons. The box plot displays the interquartile range, and the whiskers indicate the range of maximum and minimum values. The dot plots display the raw data from each individual. Statistical analyses were performed using the Wilcoxon Signed-Rank Test, with significance levels indicated as *p*<0.05 (*) and *p*<0.001 (***), *p*<0.0001 (****), while NS indicates no significant difference *p*>0.05.

## 4. Discussion

Our longitudinal survey of rubber tappers in high-risk areas of Sisaket Province identified key factors influencing anti-gSG6 P1 peptide responses of these individuals. The FAMD analysis revealed that seasonality had the strongest effect on antibody levels, demonstrating a clear correlation between seasonality and antibody responses. Among the three seasons observed, antibody levels were lowest during the rainy season, increased in the cool-dry season, and peaked in the hot-dry season. This seasonal pattern may reflect changes in human activity, as seasonal practices in rubber tapping vary. During the rainy season, the majority of participants reported entering the rubber plot 0–4 days per week (129 participants), and only 55 participants reported entering 5–7 days per week. In contrast, during the cool-dry season, the number of participants with high-frequency entry of 5–7 days/week increased to 104 participants, and that of low-frequency entry (0–4 days/week) decreased to 80 participants. In the hot-dry season, a higher proportion (99 participants) continued to enter the rubber plot frequently (5–7 days/week), while 85 participants reported lower entry frequency.

Entering rubber or forest ecotypes has been associated with increased malaria transmission risk [28]. Our findings were corroborated by an entomological survey conducted as part of a longitudinal study in Khun Han District, Sisaket Province, a historically high malaria case area [14]. The survey collected mosquitoes from two ecotypes—village and rubber-forest—to assess potential malaria transmission in each ecotype over two years, from July 2022 to March 2024, spanning the rainy (July), cool-dry (November), and hot-dry (March) seasons. Results showed that 72% of *Anopheles* mosquitoes were collected from the rubber-forest ecotype, with *An. dirus*, a major malaria vector, accounting for 84% of the total mosquitoes captured in this ecotype. *An. dirus* was present year-round, with seasonal variations in abundance. These findings suggest that participants who frequently enter the rubber plot are at higher risk of exposure to *Anopheles* mosquito bites compared to those with lower entry frequency. This pattern aligns with the antibody levels observed among participants in Sisaket Province, which frequent rubber plot entry was associated with higher antibody responses.

In addition to the spatial association, the temporal variation of the anti-gSG6 P1 peptide responses also correlate with the seasonal variation in the abundance of *Anopheles* mosquitoes. Entomological surveys in the area [14] revealed that among the 343 *Anopheles* specimens collected, the majority were from the cool-dry season (68%), followed by the rainy (24%), and the hot-dry seasons (7%). The higher number of *Anopheles* mosquitoes during the cool-dry season compared with the rainy season corresponded to a higher antibody level, increasing from 0.88 to 1.02 for an OD normalized at 410 nm. Interestingly, the antibody level remained high (1.15) during the hot-dry season, despite a low abundance of *Anopheles* mosquitoes. This suggests that the relationship between the gSG6-P1 antibody and *Anopheles* abundance is not straightforward.

One possible explanation is that the gSG6-P1 peptide used for the assay was originally developed as a marker for *Anopheles gambiae* exposure, thus appears to perform accurately in African regions where *An. gambiae* is prevalent [17, 21, 29–34]. However, studies have also observed a correlation between the antibody response against the peptide and *Anopheles* abundance in areas with limited or no *An. gambiae* presence [18, 23]. This suggests that while gSG6-P1 may serve as a general biomarker for *Anopheles* exposure, its effectiveness may vary depending on the dominant vector species in each region. Our findings are consistent with previous research indicating that the gSG6-P1 peptide may have limited sensitivity in detecting certain *Anopheles* species, such as *Anopheles arabiensis* in Tanzania [32] and *Anopheles farauti* in the Solomon Islands [35]. The variability in the specificity and sensitivity of anti-salivary peptide biomarkers across studies may be attributed to differences in sequence similarity between the *An. gambiae* salivary peptide and those of other *Anopheles* species. This discrepancy is also observed in Thailand, where the gSG6-P1 peptide was first used to assess *Anopheles* bites in Tak Province. That study found a strong correlation between *Anopheles* abundance and increasing antibody responses [23]. Unlike our study, the strong association in Tak was likely due to the predominance of *An. minimus*, whose salivary peptide shares 87% similarity with the *An. gambiae* gSG6-P1 peptide. In contrast, *An. dirus*, the dominant vector in our study area, has only 47.83% sequence similarity [23], which may explain the weaker correlation between entomological and serological data in our study. These results suggests that the *An. gambiae* gSG6-P1 peptide may not be an optimal serological tool for assessing *Anopheles* bites in regions with high species diversity such as the GMS [36–38]. To improve malaria risk assessment, there is a need to develop alternative serological biomarkers validated for different *Anopheles* species.

Several members of the Leucosphyrus group, which includes the *Anopheles leucosphyrus* and *Anopheles dirus* complexes, play a critical role in malaria transmission in the GMS. Previous ITS2 sequence alignment and phylogenetic analyses revealed a close genetic relationship among sibling species within both complexes [39, 40]. However, there is limited genetic information available on their salivary peptides. Given that different species within this group exhibit varying vector competence, it would be valuable to investigate whether salivary biomarkers could differentiate exposure to bites from vector and non-vector species. More importantly, this group of vectors in the GMS also serve as a bridge vector for zoonotic simian malaria transmission in the GMS. Developing species-specific salivary biomarkers would not only improve malaria risk assessment in humans but also provide a surrogate tool for studying *Anopheles* exposure in non-human primates. By modifying the assay to replace the secondary antibody targeting the human constant region with one targeting the simian constant region, we could assess mosquito biting exposure in monkeys. Such an assay would be a powerful tool for investigating potential reservoirs of zoonotic malaria transmission. This is particularly relevant for *Anopheles* species within the Leucophyrus group, which includes several primary malaria vectors, such as *An. dirus*, *An. hackeri*, *An. cracens*, *An. latens*, *An. introlatus*, and *An. balabacensis*— capable of transmitting both human and simian malaria parasites [41]. These species predominantly breed in forested habitats [42] and are confirmed vectors involved in the spillover of simian malaria to humans [43]. The increasing concern over zoonotic malaria infections is emphasized by reported cases in Thailand [44–47], Malaysia [48–52], and other Southeast Asian countries [53]. Therefore, the development of novel salivary peptides specific to *Anopheles* species in the Leucophyrus group would enhance our ability to assess biting exposure in both humans and monkeys. Such advancements would provide crucial insights into vector-host interactions, support malaria surveillance, and contribute to more effective control strategies for both human and zoonotic malaria transmission.

Our study demonstrates that focusing solely on *Anopheles* abundance does not fully represent the seasonal variation in *Anopheles* bite risk. Human activity plays a crucial role in determining exposure, particularly since the frequency of entering rubber-forest ecotypes varies by season, implying that the temporal risks of *Anopheles* bites has both entomological and human behavioral dimensions. During the rainy season, participants who entered rubber plots more than four days per week exhibited significantly higher antibody levels compared to those who entered less frequently. However, we observed that during the cool-dry season, when a high number of *Anopheles* mosquitoes were collected, antibody levels did not significantly differ between participants with varying frequency of rubber plot entry. This suggests that even a low frequency of entry may result in sufficient mosquito bites to elicit antibody responses when *Anopheles* density is high. Furthermore, during the hot-dry season, similar high antibody responses were observed regardless of entry frequency. However, this observation cannot be definitively claimed as there is no available data on the number of mosquito bites required to elicit an antibody response.

Another interesting aspect of our findings is that antibody responses remained high during the hot-dry season (April) despite low *Anopheles* abundance [14]. This is likely due to the carry-over effect of antibody responses from the previous season. Additionally, the entomological study only survey mosquito abundance every four months (July, November, and March) thus it is possible that *Anopheles* abundance remains high through December or January. To further investigate this, we estimated *Anopheles* abundance using Pearson’s correlation between rainfall and mosquito collections, as monthly entomological data for Sisaket Province is not available. The analyses demonstrated a delay in *Anopheles* mosquito abundance following rainfall, ranging from 2 to 4 months [14]. Given that rainfall persists until October [24], mosquito populations likely remain elevated until January or February, leading to continued biting exposure during this period. Once antibody is produced, it persist in the bloodstream for up to eight weeks (Dr. Anne Poinsignon, personal communication). Therefore, ongoing bites during January and February may sustain high antibody levels into the hot-dry season, even when *Anopheles* abundance declines. Additionally, the hot-dry season is the peak period for participants entering rubber plots, where they may receive repeated bites due to the presence of permanent *An. dirus* larval sites in these areas [14]. To gain better understanding and providing more concrete evidence of such complex relationship between the anti-salivary peptide and *Anopheles* biting risks, a more detailed longitudinal study with frequent entomological, human behavioral and serological data collection is needed. Such research will allow for a more comprehensive assessment of the temporal dynamics of antibody responses in relation to human activity and *Anopheles* abundance. Ultimately, these insights will improve our understanding of malaria transmission, particularly in the context of outdoor exposure, and will help refine vector control strategies.

Our current study highlights critical research gaps in the kinetics of anti-salivary peptide responses to *Anopheles* bites, which should be addressed for the reliable implementation of this biomarker as a survey tool. Key questions include: 1) what is the magnitude of the antibody response following mosquito bites at varying biting intensities?, 2) how quickly do antibody levels increase after exposure to mosquito bites?, 3) what is the decay rate of anti-salivary peptide antibodies over time?, 4) how does repeated exposure to mosquito bites affect the antibody response?, and 5) how do antibody responses vary after bites from different *Anopheles* species? Addressing these questions through controlled laboratory studies and subsequent field validations will provide deeper insights into the dynamics of immune responses to *Anopheles* bites and improve the utility of salivary peptide-based biomarkers for malaria risk assessment and improve vector surveillance strategies.

## Conclusions

This serological survey using an anti-gSP6-P1 salivary biomarker highlights the elevated malaria risk among individuals with a frequent entry into rubber plots, where *An. dirus* is abundant. These findings emphasize the need for targeted protective measures specifically designed to mitigate the risk posed by outdoor-biting mosquitoes. While *Anopheles* salivary peptides have shown promise in identifying high-risk groups for malaria infection, further refinement and validation are required to improve their applicability and reliability across diverse ecological and epidemiological settings, especially with the emergence of zoonotic simian malaria transmission. Strengthening serological tools for malaria risk assessment will support more effective vector surveillance and malaria control strategies in endemic regions.

## Acknowledgements

The authors sincerely appreciate the study volunteers for their time and kindness in participating in this research. We also extend our heartfelt thanks to the staff of the Ministry of Public Health for their guidance and support in facilitating fieldwork in Sisaket Province.

## Author contributions

<u>Conceptualization:</u> Sylvie Manguin, Natapong Jupatanakul

<u>Formal analysis:</u> Manop Saeung, Natapong Jupatanakul

<u>Funding acquisition:</u> Theeraphap Chareonviriyaphap, Natapong Jupatanakul, Sylvie Manguin

<u>Investigation:</u> Manop Saeung, Natapong Jupatanakul, Niramon Jampeesri

<u>Methodology:</u> Manop Saeung, Natapong Jupatanakul, Sylvie Manguin

<u>Project administration:</u> Manop Saeung, Natapong Jupatanakul, Theeraphap Chareonviriyaphap

<u>Ressources:</u> Sylvie Manguin, Theeraphap Chareonviriyaphap, Natapong Jupatanakul

<u>Software:</u> Manop Saeung, Natapong Jupatanakul, Aneta Afelt

<u>Supervision:</u> Sylvie Manguin, Theeraphap Chareonviriyaphap, Natapong Jupatanakul

<u>Validation:</u> Manop Saeung, Natapong Jupatanakul

<u>Visualisation:</u> Manop Saeung, Natapong Jupatanakul

<u>Writing –original draft:</u> Manop Saeung, Natapong Jupatanakul

<u>Writing –review & editing:</u> Manop Saeung, Sylvie Manguin, Theeraphap Chareonviriyaphap, Natapong

Jupatanakul, Niramon Jampeesri, Aneta Afelt

## Ethics approval and consent to participate

The study received ethical approval from the Research Ethics Review Committee for Research Involving Human Research Participants at Kasetsart University (Certificate of Approval No. CAO63/035). Formal written ethical clearance for the study protocol and written informed consent from volunteers were obtained before the trials commenced.

## Data Availability

All data supporting the conclusions of this article are included within the article and supporting materials.

